# Prenatal exposure to valproic acid induces sex-specific alterations in cortical and hippocampal neuronal structure and function in rats

**DOI:** 10.1101/2024.09.03.611039

**Authors:** Olivia O. F. Williams, Madeleine Coppolino, Cecilia B. Micelli, Ryan T. McCallum, Paula T. Henry-Duru, Joshua D. Manduca, Jasmin Lalonde, Melissa L. Perreault

## Abstract

**Background:** There are substantial differences in the characteristics of males and females with an autism spectrum disorder (ASD), yet there is little knowledge surrounding the mechanistic underpinnings of these differences. The valproic acid (VPA) rodent model is the most widely used model for the study of idiopathic ASD, but almost all of the studies have used male rodents.

**Method:** To fill this knowledge gap, we evaluated sex differences for neuronal activity, morphology, and glycogen synthase kinase-3 (GSK-3) signaling in primary cortical (CTX) and hippocampal (HIP) neurons prepared from rats exposed to VPA *in utero*. *In vivo*, sex-specific VPA-induced alterations in the frontal CTX transcriptome at birth were also determined.

**Results:** Overall, VPA induced more robust changes in neuronal function and structure in the CTX than in the HIP. Male- and female-derived primary CTX neurons from rats exposed to prenatal VPA had elevated activity and showed more disorganized firing. In the HIP, only the female VPA neurons showed elevated firing, while the male VPA neurons exhibited disorganized activity. Dendritic arborization of CTX neurons from VPA rats was less complex in both sexes, though this was more pronounced in the females. Conversely, both female and male HIP neurons from VPA rats showed elevated complexity distal to the soma. Female VPA CTX neurons also had an elevated number of dendritic spines. The relative activity of the α and β isoforms of GSK-3 were suppressed in both female and male VPA CTX neurons, with no changes in the HIP neurons. On postnatal day 0, alterations in CTX genes associated with neuropeptides (e.g., *penk*, *pdyn*) and receptors (e.g., *drd1*, *adora2a*) were seen in both sexes, though they were downregulated in females and upregulated in males.

**Limitations:** Primary neuron studies may not recapitulate findings performed *in vivo* or at later stages of development.

**Conclusion:** Together these findings suggest that substantial sex differences in neuronal structure and function in the VPA model may have relevance to the reported sex differences in idiopathic ASD.

## Background

Autism spectrum disorder (ASD) comprises a group of heterogeneous neurodevelopmental disorders characterized by behavioral core features that include altered social communication and repetitive stereotypic behaviors [1,2]. ASD is diagnosed up to four times more often in males than females [1–3], though it should be noted that females tend to more effectively mask their symptoms [4]. This symptom masking likely contributes to delayed or misdiagnoses in females, indicating that the reported prevalence may be erroneous [3,4]. Sex-dependent differences in etiology, age of onset, and presentation of symptoms also exist [3,5–8]. For example, it has been suggested that more males are diagnosed with “high functioning” ASD while more females are diagnosed with “low functioning” ASD [9,10]. Specifically, males display more restricted interests/repetitive behaviors, with greater difficulties in social interactions and communication [3,6,7]. At the same time, females have greater difficulty internalizing and expressing their emotions but can communicate better and are more sociable [3,6,7].

There is an overall lack of consensus of the cellular mechanisms that contribute to ASD. For example, in humans, some reports show that serum glutamate [11,12] and gamma-aminobutyric acid (GABA) [13–15] levels are elevated in autistic individuals, whereas the reduced activity of serum glycogen synthase kinase-3 (GSK-3) has been reported [16]. Conversely, other reports show evidence of reduced serum glutamate [17,18], and GABA [19], the latter of which may be age-dependent [13]. Further, GSK-3 inhibitors have been suggested to have therapeutic potential, implying elevated GSK-3 activity [20,21]. In animal models used to study various aspects of ASD, there appear to be significant variations in the findings. For example, studies examining alterations in neuronal morphology in these models are inconsistent, with differential findings concerning neuronal dendrite length, dendritic spine number, complexity, and soma volume [16,22–26]. Alterations in GSK-3 activity have also been described, with the kinase being overactive [27] or underactive [16,28] in different models of ASD. Overall, these clinical and preclinical discrepancies are most likely explained by the heterogeneity of ASD, variations in sex and age of the subjects studied [5,29], and the various genetic or idiopathic models used to study this group of disorders.

The lack of consensus in many reported findings highlights the need to be cautious when making broad assumptions about ASD, and emphasizes extreme caution when pooling subjects [30]. Not only are age and sex key variables in ASD, but consideration of how the disorder manifests at both a behavioral and emotional level is paramount, as individual differences in the disorder phenotype almost certainly reflect differences in the mechanistic underpinnings. An excellent example are studies examining neuronal systems function, a process known to be highly coupled to behavioral states [31] and that has been suggested to be a potential biomarker in ASD [32]. However, it must be emphasized that any identified biomarker would be relevant to only a subset of those that have ASD, based on comorbidities and symptoms, but also would vary with age and sex [29]. Indeed, preclinical work has shown that changes in neuronal oscillatory activity in response to stress can vary between sexes, despite the accompanying behaviors looking similar [33], indicating that male- and female-specific biomarkers would most likely need to be identified whether or not sex differences in behavior were evident.

Preclinical studies that use models of genetic or idiopathic ASD have historically shown a lack of consideration of sex as an experimental variable that has likely had long-term impacts on translatability. For example, the most widely used model of idiopathic ASD is the valproic acid (VPA) rodent model, which displays face, construct, and predictive validity [34]. Of the hundreds of studies using this model, only a handful of studies have examined sex differences, with behavioral studies showing that VPA-exposed females are more sociable than males [35–37], one of the core symptoms of the disorder. As the VPA model of ASD is extensively used, this study sought to provide an in-depth characterization of sex differences in neuronal structure and function in postnatal neurodevelopment using rat primary cortical (CTX) and hippocampal (HIP) neurons derived from rat offspring exposed *in utero* to VPA. VPA-induced sex-specific alterations in frontal cortical gene expression *in vivo* on postnatal day 0 were also characterized.

## Methods

### Animals

Eighteen pregnant Sprague-Dawley dams (Charles River, QC) and their offspring were used. Dams were individually housed in large opaque rodent cages with standard enrichment and kept on a 12-hour reverse light-dark cycle, food, and water *ad libitum*. The VPA model of idiopathic ASD was used [34]. It must be noted that this model system represents a single dimension of the disorders that exist along the spectrum, though may have relevance to several other environmental drug and/or toxin exposures *in utero* [34]. Pregnant dams were administered a single intraperitoneal (i.p.) injection of VPA (500 mg/kg, concentration 250 mg/mL; Sigma-Aldrich Canada) or saline (SAL) (1 mL/kg) on gestational day 12.5 and left undisturbed until parturition. All protocols were under the guidelines set out by the Canadian Council Animal Care Committee at the University of Guelph.

### Primary Neuronal Cell Cultures

Primary neuronal CTX and HIP cultures derived selectively from female and male offspring were generated from postnatal day 0-1 rat pups as previously described [38]. The offspring were sexed by anogenital distance and divided into four experimental groups: female saline (FSAL), male saline (MSAL), female VPA (FVPA), and male VPA (MVPA). Following rapid decapitation CTX and HIP tissue were dissected and placed in dissection media (water, 1x HBSS, 1% 1 M HEPES, 1% penicillin/streptomycin (Gibco; Fisher Scientific, Mississauga, ON, Canada)) on ice. The collected tissue was washed and dissociated in 0.5% Trypsin-EDTA (Gibco) at 37°C for 20 minutes. Following washing with dissection media, tissue was triturated, passed through a 100 μm pore cell strainer (Fisher Scientific, Ottawa, ON, Canada), and centrifuged at 200 x *g* for 5 minutes at room temperature. The supernatant was aspirated, and cells were resuspended in plating media (Neurobasal medium, 2% B27 supplement, 1% 200 mM L-glutamine, 1% penicillin/streptomycin (Gibco), 5% Fetal bovine serum (Cytiva, Fisher Scientific, Ottawa, ON, Canada)). Cells were counted (Bio-Rad cell counter, Mississauga, ON, Canada) with trypan blue (Bio-Rad, Mississauga, ON, Canada) and diluted with plating media to a density of 1 x 10^6^ cells/mL. Cells were plated at a density of 3 x 10^5^ cells/well on 24-well CytoView Multi-electrode array (MEA) plates (Axion Biosystems, Atlanta, GA, USA) for electrophysiological recordings; 2 x 10^5^ cells/well on 24-well Corning plates with 12 mm glass coverslips (Fisher Scientific) for immunocytochemistry; and 1 x 10^6^ cells/well on 6-well Corning plates for western blot analysis. All plates were coated with 0.1 mg/mL poly-D-lysine for CTX cells and 0.1 mg/mL poly-L-lysine for HIP cells with 2.5 μg/mL laminin (SigmaAldrich). Cell cultures underwent half-media changes with serum-free media (Neurobasal Medium, 2% B27 supplement, 1% 200 mM L-glutamine, and 1% penicillin/streptomycin; all Gibco) two hours post-plating and maintained every 3-4 days for a total culturing period of 21 days. On days *in vitro* (DIV) 1, cultures grown for immunocytochemistry or western blot analysis received a single treatment of cytosine Beta-D-arabinofuranoside (AraC) (10 µM; Sigma-Aldrich), as AraC reduces proliferation of non-neuronal cells, resulting in neuron-only cultures [39]. Cultures grown for MEA analysis did not receive AraC, as non-neuronal cells within the cultures promote the formation of electrically active neuronal networks. MEA recordings, immunocytochemistry and protein extraction were performed on DIV21.

### Multielectrode array recordings

MEA recordings are used to measure the electrical activity of groups of neurons grown *in vitro*, and provide insights into the neural dynamics of a population of cells [40]. MEA recordings were performed from the cultures on DIV21 using the Maestro Edge multi-well MEA recorder (Axion Biosystems, Atlanta, GA, USA). Axion’s Integrated Studio software (AxIS, Axion Biosystems, Atlanta, GA, USA) was used to record spontaneous neuronal activity [38]. MEA recordings were 30 minutes long and taken at a sampling frequency of 12.5 kHz. Files were processed and analyzed using the NeuralMetric Tool software (Axion Biosystems). Throughout the recording process, plates were maintained at 37°C and active electrodes were defined as >5 spikes per minute. The mean firing rate (MFR), number of bursts, number of spikes, inter-spike interval (ISI) within a burst, and mean number of spikes/burst were extracted using data from all active electrodes within a treatment group. The synchrony index and coefficient of variation (CV) for ISI were calculated for each well with more than 5 active electrodes. Following spontaneous recording, extracellular waveform data were exported and analyzed with the Plexon Offline Sorter spike sorting tool (Offline Sorter v4.7.1, Plexon Inc., Dallas, TX, USA). Average peak, valley, and peak-valley distance were measured from each extracellular spike waveform that occurred over the 30 minute recording on each active electrode.

### Immunocytochemistry

Immunocytochemistry (ICC) was performed as previously described [41]. 500 µL of 8% paraformaldehyde (PFA) was added to 500 µL of media to obtain a working concentration of 4% PFA. PFA was added to each well, and cells were incubated for 30 minutes at room temperature. Cells were washed three times in cell culture grade phosphate-buffered saline (PBS) (Gibco) for 10 minutes and blocked in 4% bovine serum albumin (BSA) (Thermo Fisher Scientific, Canada), 0.1% Triton X-100 (Fisher Scientific, Ottawa, ON, Canada), and 5% normal goat serum (NGS) in 1x PBS for 60 minutes at room temperature. Cells were immunostained with MAP2 primary antibody (1:500, rabbit, ab5622, Millipore, Sigma-Aldrich, Oakville, ON, Canada), β-Actin (1:500, mouse, ab8226, Abcam), Drebrin A (1:500, rabbit, #12243, Cell Signaling) and/or Ankyrin G (1:500, mouse, #75-146 Antibodies Inc), followed by incubation with the appropriate anti-rabbit or anti-mouse secondary antibody (Alexa Fluor 488 or 594, Invitrogen). Slides were coverslipped with Prolong Gold (Invitrogen), kept overnight at 4°C, and subsequently imaged with an Olympus FV1200 Confocal Microscope. Images were taken at 20X or 60X magnification.

### Sholl Analysis

To analyze dendritic arborization, image stacks were taken with a step size in the Z-axis of 1 μm using 20X magnification. Neurons were reconstructed with Neuromantic and Sholl analysis performed using ImageJ software (https://imagej.nih.gov/ij/download.html). Traces were used to evaluate the mean number of dendritic intersections, measured between concentric rings radiating from the soma in 10 μm increments. The neuron tracings were also used to measure differences in the average of the longest dendrite of the neurons between groups.

### Western blotting

Western blotting was performed as previously described [42]. Cells were washed twice with ice-cold 1x PBS and collected in ice-cold RIPA buffer (Sigma-Aldrich, Oakville, ON, Canada) with added phosphatase (10 μL/mL; Fisher Scientific, Ottawa, ON, Canada) and protease inhibitors (10 μL/mL; Sigma-Aldrich, Oakville, ON, Canada). Protein concentrations were determined using the Bradford assay (Biospectrometer, ThermoFisher). Protein (20 μg) was loaded onto a 10% sodium dodecyl sulfate (SDS) polyacrylamide gel for electrophoresis at 80 V for 20 minutes, and 120 V for 45 minutes using a Mini-PROTEAN Tetra cell system (Bio-Rad, Mississauga, ON, Canada). Proteins were then transferred at constant 25 V for 30 minutes onto a polyvinylidene difluoride (PVDF) membrane. Membranes were briefly washed in Tris-buffered saline (TBS) with 0.1% Tween-20 (TBS-T) and blocked in 5% non-fat milk in TBS-T and incubated overnight at 4°C in primary antibodies specific for either GSK-3 α/β, or phospho-GSK-3 α/β (GSK-3 α/β, 1:10,000, rabbit-#5676, Cell Signaling; phospho-GSK-3 α/β (Ser21/9), 1:10,000, rabbit-# 9331S, Cell Signaling). Blots were washed three times with TBS, incubated with goat anti-rabbit horseradish peroxidase (HRP)-conjugated polyclonal secondary antibody (1:10,000, Bio-Rad) for 1 hour at room temperature and washed three times with TBS for 10 minutes each. Bands were visualized using enhanced chemiluminescence (ECL) on a ChemiDoc MP imaging system (Bio-Rad, Mississauga, ON, Canada). Total protein was used as a loading control. Optical density was measured by normalizing the relative expression of each antibody to the total protein present in each lane.

### Gene Expression

Postnatal day 0 pups were sexed and euthanized via rapid decapitation, and CTX tissue was dissected and flash frozen. As previously described, total RNA was isolated using the RNeasy Mini Kit (Qiagen, Hilden, Germany) [43]. RNA quality was assessed by NanoDrop ND-1000 (260/280 >2; ThermoScientific) and by RNA ScreenTape (RIN ≥7; Agilent). Samples were then sent to the TCAG Microarray Facility at Sick Kids Hospital (Toronto) for microarray processing, which interrogates > 27,000 protein coding transcripts, with a median of 22 probes per gene, and reproducibility at a signal correlation coefficient > 0.99. All microarray data are accessible (GEO Accession # GSE275787). Analysis of genes that were transcriptionally altered was performed on GeneSpring GX 14.9.1 (Agilent Technologies Inc). Raw CEL files were uploaded into a project file with exon analysis and Affymetrix exon expression and biological significance workflow analysis. The rat gene 2.0 ST annotation technology (RaGene-2_0-st-na33_2_rn4_2013-03-28) was used for all analyses. Raw fluorescence data were normalized across all chips in each study, with a lower threshold of > 60 Raw Fluorescence Units (RFU) in at least 50% of conditions. Principal Components Analysis (PCA) was used for group-level clustering. A fold change of 1.3 was deemed the threshold for significant change to assess differences between VPA and control groups, for each sex. Gene ontology (GO) analysis for enriched genes was performed using the NIH DAVID Bioinformatics Database, Functional Annotation Tool.

### Statistical analyses

Before all analyses, Shapiro-Wilk or Kolmogorov-Smirnov normality tests were conducted depending on sample sizes (Shapiro-Wilk for N < 50, Kolmogorov–Smirnov N ≥ 50) and variance assessed by Levene’s test. Data were analyzed using a two-way analysis of variance (ANOVA) with Sex and Treatment as between-subject factors. MEA data was collected from 16 electrodes/well for 6 wells from 3 biological replicates, for a total of N = 288 electrodes/group, electrodes were excluded if activity was <5 spikes per minute (N = 136-288). Morphology was performed by assessing four neurons from 3 biological replicates for a total N = 12 neurons/group. Sholl analysis was analyzed using repeated measures ANOVA with Distance as the within-subjects factor and Treatment as the between-subjects factor. The assumption of sphericity was tested with the Mauchly’s Test, and data was analyzed separately for males and females within the CTX or HIP. Western blot analysis was performed on 6 biological replicates per group. Transcriptomics data was extracted from 5 animals per group. For all ANOVAs, group differences were assessed using Tukey or Games-Howell *post-hoc* tests as appropriate. Within-sex planned comparisons were performed between Treatments in some instances, as noted, in the absence of a Sex x Treatment interaction. All graphical data are presented as means ± SEM. Statistical significance is defined at *p* < 0.05. For the gene expression analysis, a corrected (Bonferroni) *p*-value of ≤ 0.010 was used. Statistical analyses were performed in IBM Statistical Package for the Social Sciences (SPSS, Version 28).

## Results

### VPA-derived neurons show elevated activity with enhanced effect in females

The sex-specific changes in neuronal activity of SAL- or VPA-derived neurons, *in vitro,* were measured on DIV21 from CTX and HIP primary neuronal cell cultures. F-statistics from the two-way ANOVAs for Figures 1-3 are shown in Supplemental Table 1. Representative raster plots from CTX neurons for each group across a 30 minute recording are shown (Fig. 1A), as well as a representative sample of bursting activity across one second (Fig. 1B).

**Figure 1.**
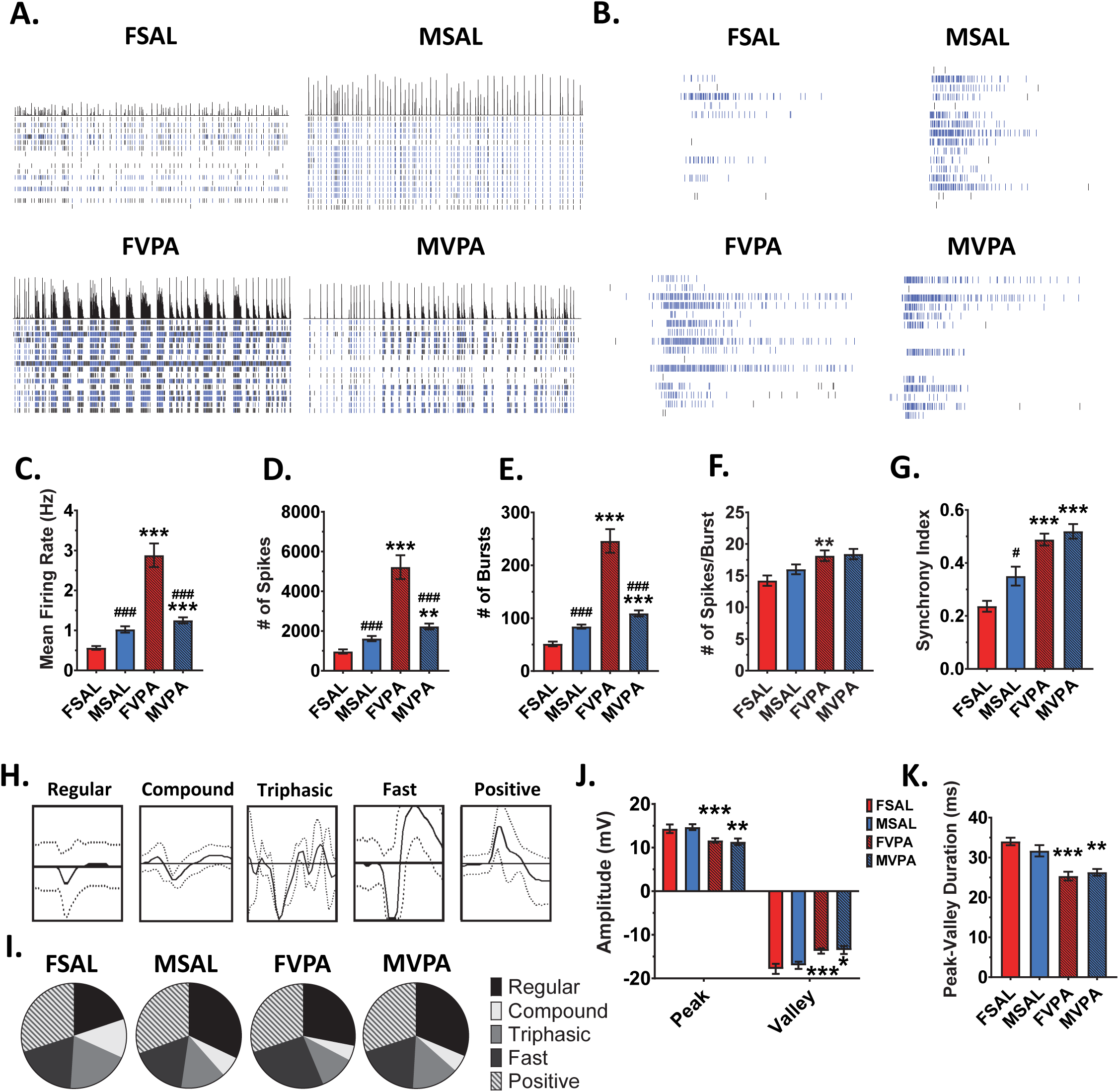
Sex-specific effect of prenatal VPA exposure on neuronal activity of CTX neurons. **A)** Raster plots for the CTX neurons MEA recording over the 30 minutes displaying spike trains at each of the twelve electrodes. **B)** Spike trains at each of the twelve electrodes within a burst for each experimental group over a 1 second bin **C-D)** VPA elevated the mean firing rate, total number of spikes and bursts. **F)** Elevated number of spikes per burst in FVPA neurons. **G)** Elevated synchrony index for both FVPA and MVPA. **H)** Representative waveforms. **I)** Proportion of waveform types across the four experimental conditions. **J)** Amplitude of the peak and valley of waveforms were reduced for both FVPA and MVPA neurons. **K)** Duration between peak and valley was reduced for FVPA and MVPA neurons. Data are expressed as means ± SEM. N = 6 wells, 3 biological replicates. * *p* < 0.05, ** *p* < 0.01, *** *p* < 0.001 compared to sex-matched controls, ^###^ *p* < 0.001 compared to females of the same model, ANOVA followed by Bonferroni or Games–Howell post-hoc determined by Levene’s test of variance.

In the CTX neurons, FVPA derived neurons showed significantly higher activity compared to sex-matched controls, as evidenced by a higher MFR (*p* < 0.001), number of spikes (*p* < 0.001), number of bursts (*p* < 0.001), and spikes/burst (*p* = 0.005) (Fig. 1C-F). CTX MVPA neurons also showed elevated activity compared to sex-matched controls with a higher MFR (*p* = 0.001), number of spikes (*p* = 0.010), and number of bursts (*p* < 0.001) (Fig.1C-E). Sex differences in CTX neuronal activity were evident between the VPA-treated groups, with the FVPA neurons showing a greater MFR (*p* < 0.001), number of spikes (*p* < 0.001), and number of bursts (*p* < 0.001) compared to MVPA neurons (Fig. 1C-E). There was a significant effect of Sex and Treatment on synchrony, such that VPA-derived neurons showed greater synchrony than SAL- derived neurons (female: *p* < 0.001, male: *p* < 0.001) (Fig. 1G). Innate sex differences between FSAL and MSAL CTX neurons were also observed, such that MSAL neurons displayed a higher MFR (*p* < 0.001), number of spikes (*p* < 0.001), number of bursts (*p* < 0.001), and synchrony (*p* = 0.028) compared to FSAL neurons (Fig. 1C-E, G). Conversely, in the VPA model, some of these inherent sex differences were abolished (synchrony) (Fig. 1G) or reversed (MFR, # of spikes, # of bursts) (Fig. 1C-E).

Extracellular waveforms can be characterized into five types: regular, compound, triphasic, fast, and positive, with representative tracings (Fig. 1H). Pie charts depicting the proportion of waveforms within each group are also shown (Fig. 1I). No significant differences were observed between groups for the proportion of waveform types. In both FVPA and MVPA CTX neurons, the amplitude of the peak was smaller than their sex-matched controls (female: *p* < 0.001, male: *p* = 0.005), as was the amplitude of valley (female: *p* < 0.001, male: *p* = 0.02) (Fig. 1J) (peak: Treatment, F(1,66) = 32.4, *p* < 0.001; valley: Treatment, F(1,66) = 38.0, *p* < 0.001). Furthermore, the duration of time between the amplitude of the peak to valley showed that the VPA neurons were hyperpolarizing at a faster rate (female: *p* < 0.001, male: *p* = 0.005) (Fig. 1K) (Treatment, F(1,66 = 40.4, *p* < 0.001).

Representative raster plots from HIP neurons for each group across a 30 minute recording are shown, as well as a representative sample of bursting activity across one second (Fig. 2A, B). In the HIP neurons, the FVPA group exhibited significantly greater activity indicated by elevated MFR (*p* < 0.001), number of spikes (*p* < 0.001), and number of bursts (*p* < 0.001) compared to the FSAL group (Fig. 2C-E). MVPA neurons had a lower number of bursts (*p* < 0.001), and a significant elevation in the number of spikes per burst (*p* = 0.006) compared to the MSAL group (Fig. 2E, F). When sex differences in the VPA-treated neurons were evaluated, the FVPA neurons were higher than the MVPA neurons in number of bursts (*p* = 0.007), while the MVPA were significantly higher than FVPA in number of spikes/burst (*p* = 0.047) (Fig. 2E, F). Comparing the control groups, MSAL neurons had elevated MFR (*p* < 0.001), number of spikes (*p* = 0.014), number of bursts (*p* < 0.001), and synchrony (*p* < 0.001) compared to FSAL neurons (Fig. 2C-E, G). Similar to the CTX, in the VPA model, many of these inherent sex differences were abolished (MFR, # of spikes, synchrony) (Fig. 2C, D, and G) or reversed (# of bursts, # of spikes/burst) (Fig. 2E, F).

**Figure 2.**
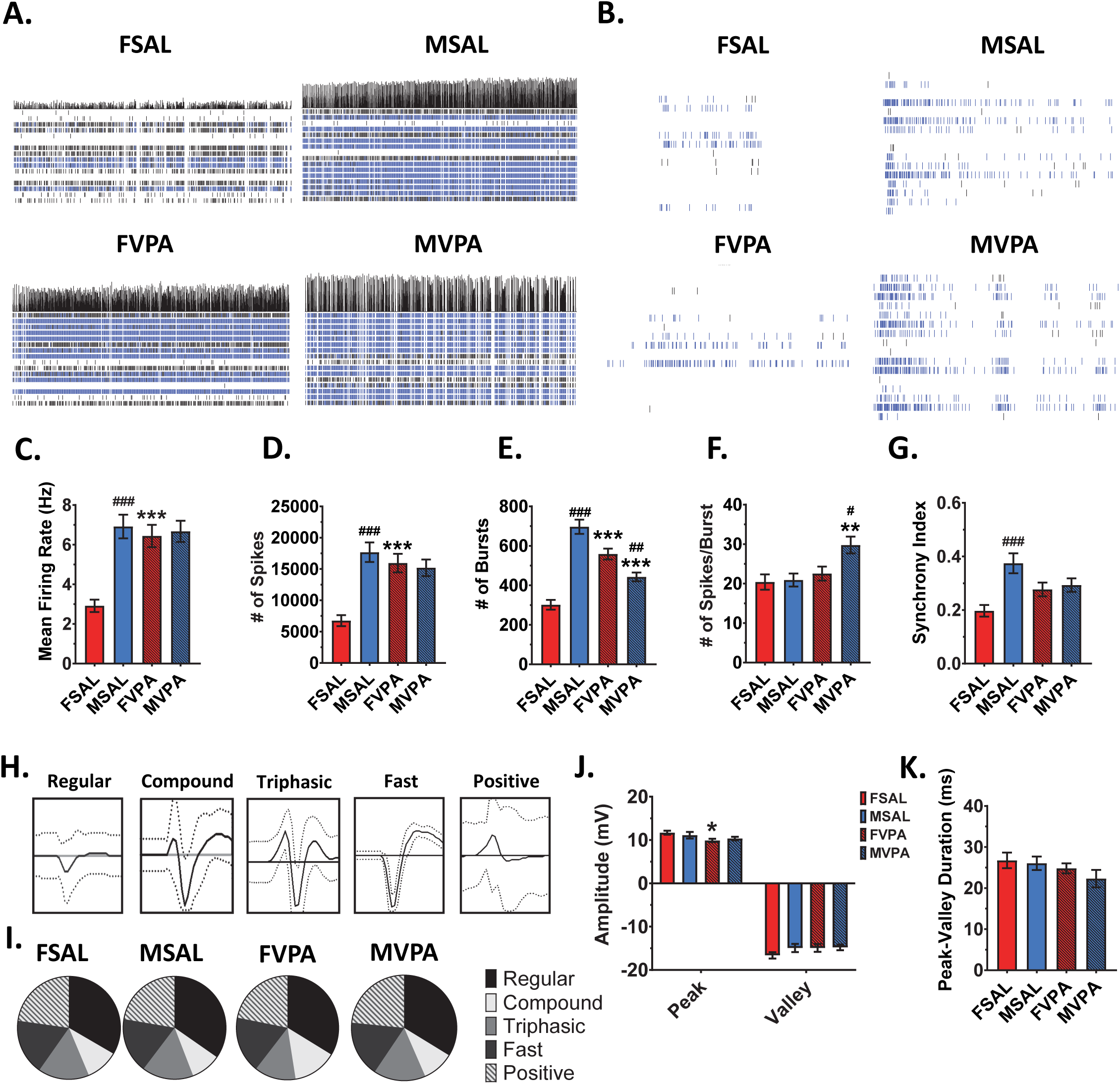
Sex-specific effect of prenatal VPA exposure on neuronal activity of HIP neurons. **A)** Raster plots for the HIP neurons MEA recording over the 30 minutes displaying spike trains at each of the twelve electrodes. **B)** Spike trains at each of the twelve electrodes within a burst for each experimental group over a 1 second bin. **C, D)** Elevated mean firing rate and number of spikes in FVPA neurons. **E)** Elevated number of bursts in FVPA, but reduced number of bursts in MVPA neurons. **F)** Elevated number of spikes per burst in MVPA neurons. **G)** No significant effect of VPA on the synchrony index. **H)** Representative waveforms. **I)** Proportion of waveform types across the four experimental conditions. **J)** Reduced peak amplitude in FVPA neurons. **K)** No significant difference in the duration of peak to valley. Data are expressed as means ± SEM. N = 6 wells, 3 biological replicates. * *p* < 0.05, ** *p* < 0.01, *** *p* < 0.001 compared to sex-matched controls, ^#^ *p* < 0.05, ^##^ *p* < 0.01, ^###^ *p* < 0.001 compared to females of the same model, ANOVA followed by Bonferroni or Games–Howell post-hoc determined by Levene’s test of variance.

Representative tracings of the five types of waveforms found in HIP recordings are shown (Fig. 2H). Pie charts depicting the proportion of each waveform for the four treatment groups are displayed (Fig. 2I). No significant differences were observed between groups for the proportion of waveform types. Unlike the CTX, only the amplitude of the peak was smaller for the FVPA neurons compared to sex-matched controls (*p* = 0.027) (Fig. 2J) (Treatment, F(1,65)=5.8, *p* = 0.018). No significant differences were observed in the amplitude of the valley in HIP or in the distance between the peak and valley amplitudes in either sex (Fig. 2J, K).

For the organization of neuronal firing, the relationship between ISI and the number of spikes was examined for each group, with representative correlograms for CTX and HIP shown (Fig. 3A, D). In the CTX, the ISI was lower for both female and male VPA-derived neurons (female: *p* < 0.001, male: *p* < 0.001) (Fig. 3B). Neuronal spike variability, or predictability of firing, can be extrapolated from the correlograms and is represented as the CV. Both groups of VPA-derived neurons had elevated CV (female: *p* < 0.001, male: *p* < 0.001), revealing a disorganization in firing patterns (Fig. 3C). In the HIP, there was a significant but modest suppression of ISI in the FVPA neurons (*p* = 0.047) and an elevation in MVPA neurons (*p* < 0.001) compared to sex-matched controls (Fig. 3E). Similarly, compared to controls, FVPA neurons had a lower CV (*p* < 0.001), while the CV of MVPA neurons was higher (*p* < 0.001) (Fig. 3F). These findings indicate that VPA-induced disorganization of firing in the HIP was male-specific.

**Figure 3.**
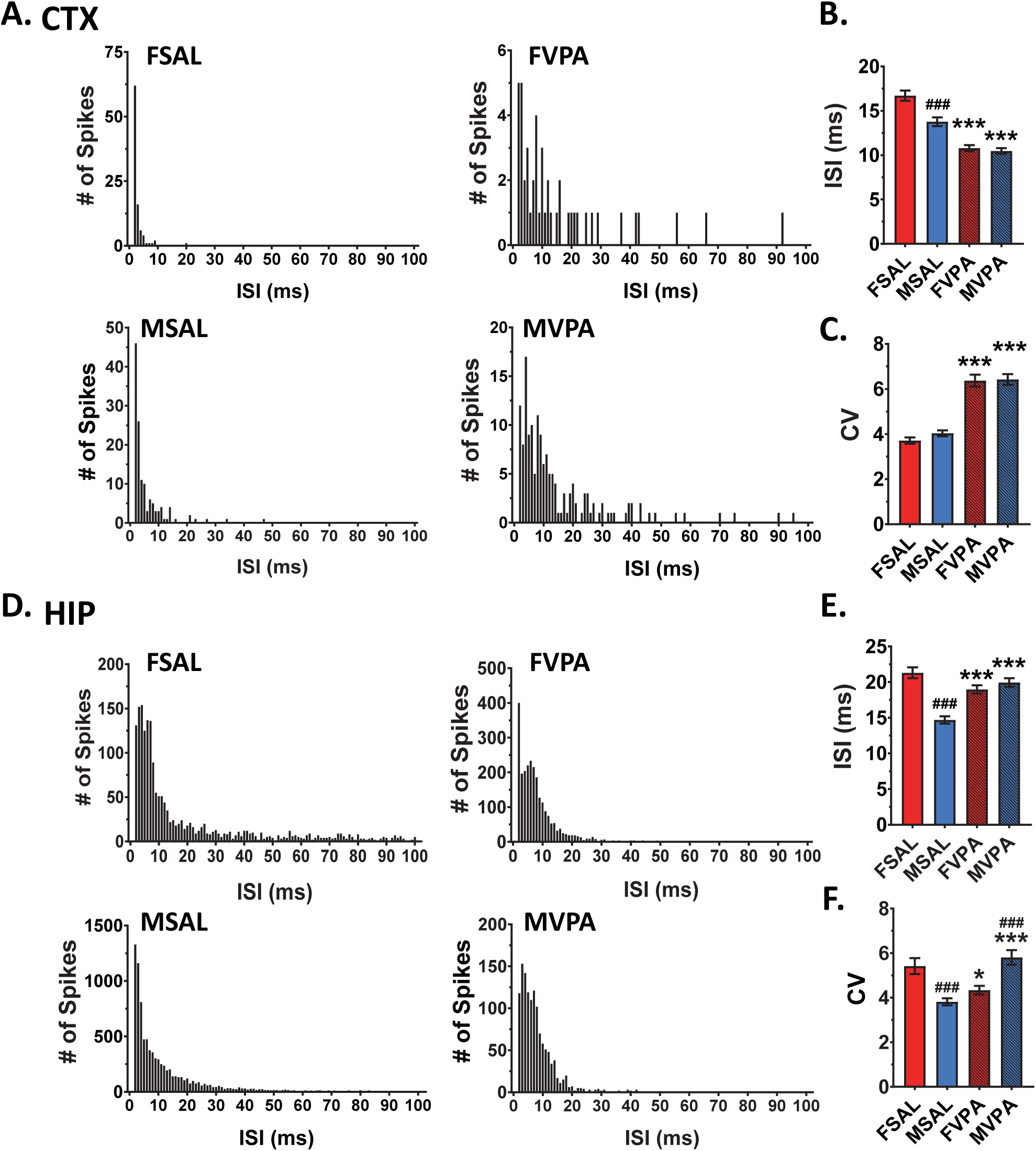
VPA alters organizational firing in CTX and HIP neurons. **A)** Representative ISI histograms for the four experimental conditions in the CTX. **B)** Reduced ISI, within a burst, was evident in both VPA groups. **C)** Elevated CV in both FVPA and MVPA CTX neurons indicates disorganized firing. **D)** Representative ISI histograms for the four experimental conditions in the HIP. **E)** MVPA neurons have a higher ISI within a burst. **F)** CV was elevated for MVPA HIP neurons, but reduced for FVPA neurons, suggesting MVPA neurons have more disorganized firing, while FVPA neurons have more predictable firing patterns. Data are expressed as means± SEM. N = 6 wells, 3 biological replicates. * *p* < 0.05, *** *p* < 0.001 compared to sex-matched controls, ^###^ *p* < 0.001 compared to females of the same model, ANOVA followed by Bonferroni or Games–Howell post-hoc determined by Levene’s test of variance.

### VPA neurons show sex- and region-specific alterations in complexity

We next sought to evaluate whether there were sex and/or treatment differences in neuronal morphology. Representative images of CTX neurons on DIV21 showing dendritic complexity are shown in Figure 4A. Representative group images depicting the axon initial segment (AIS), the site of action potential initiation, of CTX neurons are shown in Figure 4B.

**Figure 4.**
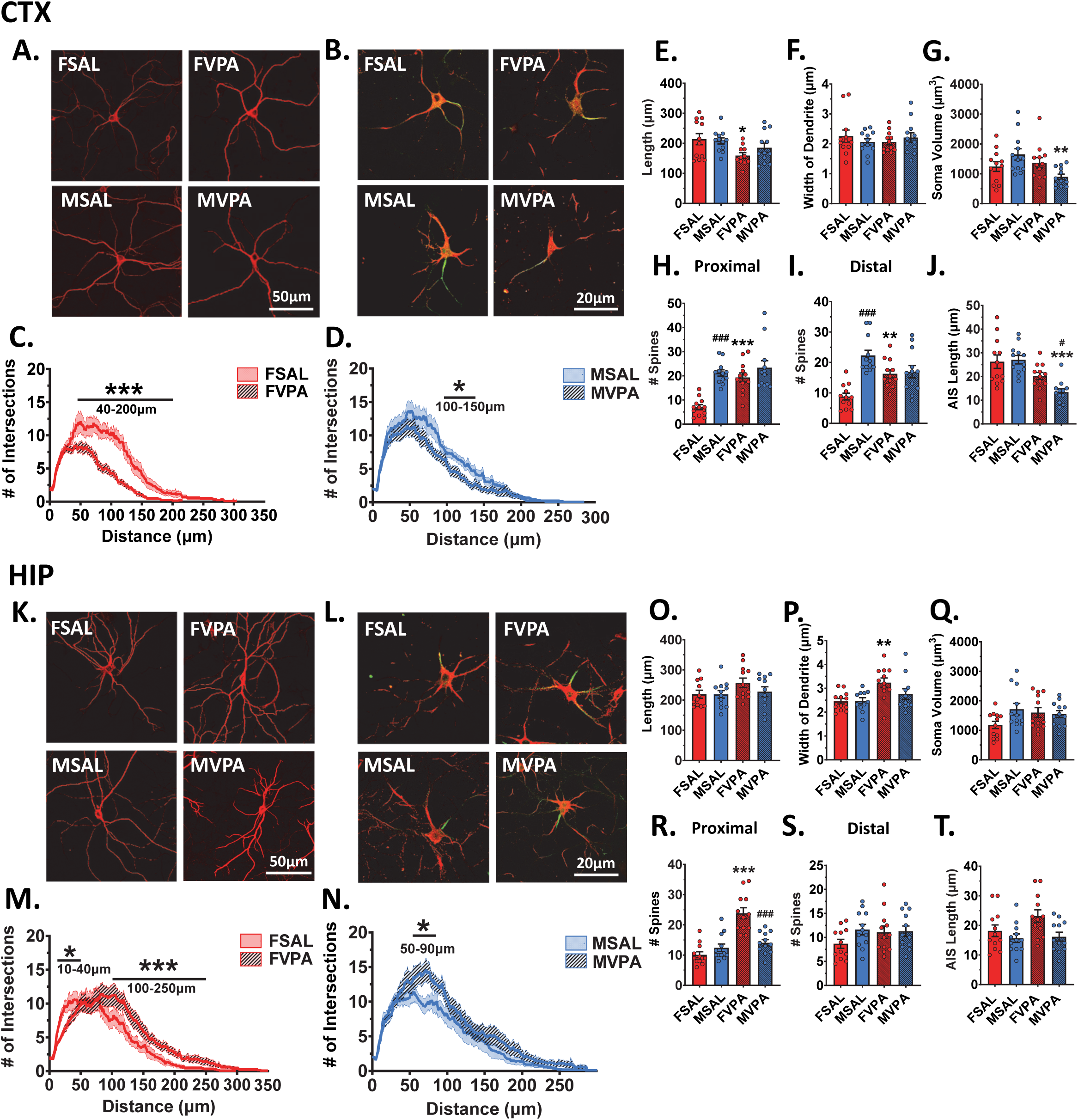
Sex-specific VPA-induced alterations in dendritic morphology. **A)** Representative zoomed images for CTX neurons for the four experimental conditions imaged at 20X magnification. **B)** Representative CTX neurons showing the AIS imaged at 60X magnification**. C, D)** Sholl analysis showed reduced dendritic complexity in FVPA and MVPA neurons compared to sex-matched controls. **E)** FVPA neurons have shorter dendrite length. **F)** No differences in the width of the neurons were evident between groups. **G)** MVPA neurons had reduced soma volume. **H, I)** FVPA neurons had more dendritic spines proximal (<30um) and distal (>30um) to the soma. **J)** MVPA neurons had a shorter AIS compared to controls. **K)** Representative zoomed images of HIP neurons for the four experimental conditions imaged at 20X magnification. **L)** Representative HIP neurons showing the AIS imaged at 60X magnification. **M)** Sholl analysis for the female neurons, FVPA neurons have an initial decrease followed by increased dendritic complexity. **N)** Sholl analysis for male neurons, MVPA neurons have a moderate increase in complexity. **O)** No significant differences in dendrite length, **P)** FVPA neurons had wider dendrites compared to sex-matched controls. **Q)** No significant difference in soma volume between groups. **R)** FVPA neurons had more dendritic spines proximal (<30 µm) to the soma compared to controls. **S)** No significant group differences in dendritic spine number distal (>30um) to the soma. **T)** No significant group differences in AIS length. Microtubule-associated protein 2 antibody (MAP2) and ankyrin G antibodies were used for representative images. Data are expressed as means ± SEM. N = 3 biological replicates with 4 neurons per replicate * *p* < 0.05, ** *p* < 0.01, *** *p* < 0.001 compared to sex-matched controls, ^#^ *p* < 0.05, ^##^ *p* < 0.01, ^###^ *p* < 0.001 compared to females of the same model, ANOVA followed by Bonferroni or Games–Howell post-hoc determined by Levene’s test of variance. Within-subject effects for the Sholl analyses were evaluated by repeated measures ANOVA with Treatment as the between-subject factors.

A repeated measures ANOVA assessing group differences in the number of dendritic intersections per unit distance revealed a significant effect of Treatment and a Sex by Treatment interaction (Figs. 4C, D; Treatment, F(1,44 = 16.1, *p* < 0.001; Sex by Treatment, F(1,44) = 4.2, *p* = 0.046), with FVPA rats showing reduced CTX dendritic complexity compared to sex-matched controls (*p* < 0.001) and no differences between the male groups. To more closely examine VPA-induced differences in dendritic complexity within sex, comparisons were performed at select distances from the soma. In female-derived CTX neurons, there were fewer intersections in neurons from the FVPA group at 40-200 μm from the soma (Fig. 4C) (F(1,22) = 17.7 *p* < 0.001). A similar loss of complexity was evident in the MVPA neurons, though in a much narrower range, 100-150 μm from the cell soma (Fig. 4D; F(1,22) = 6.8, *p* = 0.016).

Dendritic length, soma volume, proximal dendrite width, dendritic spines (proximal and distal), and AIS were next assessed. FVPA, but not MVPA, CTX neurons showed reduced overall dendrite length compared to sex-matched control neurons (*p* = 0.029) (Fig. 4E), with no group differences in dendritic width (Fig. 4F). Conversely, only MVPA neurons showed a reduced soma volume compared to MSAL neurons (*p* = 0.009) (Fig. 4G) (Dendrite length: Treatment, F(1,41) = 11.1, *p* = 0.002; Soma Volume: Treatment, F(1,44) = 3.8, *p* = 0.05; Sex by Treatment, F(1,44) = 7.6, *p* = 0.008). FVPA neurons had more dendritic spines compared to FSAL neurons both proximally (<30 µm, *p* < 0.001) and distally (>60 µm, *p* = 0.009) to the soma (Fig. 4H, I). An inherent sex difference was observed wherein there were more dendritic spines on MSAL neurons compared to FSAL, both proximally (*p* < 0.001) and distally (*p* < 0.001), a sex-difference not evident between the VPA groups (Fig. 4H, I) (Proximal dendritic spines: Sex, F(1,44 = 22.9, *p* < 0.001; Treatment, F(1,44) = 14.7, *p* < 0.001; Sex by Treatment, F(1,44) = 7.1, *p* = 0.011; Distal dendritic spines: Sex, F(1,44) = 19.8, *p* < 0.001; Sex by Treatment, F(1,44) = 16.3, *p* < 0.001). AIS length was lower for MVPA (*p* < 0.001) CTX neurons compared to sex-matched controls (Fig. 4J). A sex difference was also observed in the VPA neurons, in which the MVPA AIS was significantly shorter than the FVPA AIS (*p* = 0.016) (Fig. 4J) (Treatment, F(1,44) = 25.4, *p* < 0.001; Sex by Treatment, F(1,44) = 3.7, *p* = 0.05).

Dendritic morphology was also evaluated in the HIP with representative images shown in Figure 4K, and representative group images showing the AIS of HIP neurons in Figure 4L. VPA induced significant distance-dependent changes in HIP dendritic complexity in the FVPA and MVPA neurons (Figs. 4M, N). A repeated measures ANOVA assessing the group differences in the number of dendritic intersections per unit distance (Fig. 4M, N) revealed a significant effect of Treatment (Treatment, F(1,38 = 7.6, *p* = 0.009) though no significant group differences were observed. Within sex comparisons at select distances from the soma showed that in FVPA HIP neurons, the number of intersections were decreased at 10-40 µm from the soma, and increased 100-250 µm from the soma compared to controls (Fig. 4M) (10-40 µm, F(1,22) = 5.9, *p* = 0.026; 100-250 µm, F(1,19) = 16.7, *p* < 0.001). MVPA neurons showed increased complexity 50-90µm from the soma (Fig. 4N) (F(1,19) = 6.7, *p* = 0.018).

There were no significant group differences in HIP neuron length or soma volume (Fig. 4O, Q). In the FVPA neurons, proximal dendritic width was greater compared to their controls (*p* = 0.01) with no differences between male groups (Fig. 4P) (Treatment, F(1,42) = 9.9, *p* = 0.003). FVPA neurons had a greater number of dendrites compared to controls (*p* < 0.001), with no differences between the male groups (Fig. 4R) (Sex, F(1,44) = 8.0, *p* = 0.007; Treatment, F(1,44) = 34.1, *p* < 0.001; Sex by Treatment, F(1,44) = 20.9, *p* < 0.001). There was a main effect of Sex when AIS length was examined, with female-derived neurons having an overall longer AIS than the male-derived neurons (Fig. 4T) (Sex, F(1,44) = 7, *p* = 0.011).

### VPA induces sex- and region-specific alterations in GSK-3 signaling

The sex- and region-dependent effects of VPA on GSK-3 expression and activity were next evaluated. Representative blots displaying expression levels of phosphorylated and total GSK-3 α and β are shown for the CTX and HIP (Fig. 5A, B). FVPA CTX neurons had lower overall levels of pGSK-3α (*p* = 0.049) and pGSK-3β (*p* < 0.001) compared to FSAL neurons, whereas MVPA CTX neurons had elevated levels of pGSK-3α (*p* = 0.003) and pGSK-3β (*p* = 0.01) compared to their sex-matched controls (Fig. 5C, left panel). In the control neurons, MSAL neurons showed innately lower pGSK-3α expression (*p* = 0.013) compared to FSAL neurons (Fig. 5C, left panel; α: Sex, F(1,20) = 8.4, *p* = 0.009; Treatment, F(1,20) = 17.3, *p* < 0.001; Sex by Treatment F(1,20) = 58.3 *p* < 0.001; β: Sex, F(1,20) = 4.6, *p* = 0.045; Sex by Treatment, F(1,20) = 34.5, *p* < 0.001). Concerning total CTX GSK-3 expression (Fig. 5C, center panel), FVPA and MVPA neurons had lower GSK-3α expression compared to sex-matched controls (female: *p* < 0.001; male: *p* = 0.021), whereas for GSK-3β only FVPA neurons showed this reduction (*p* < 0.001). No sex differences in total GSK-3 expression were observed between the SAL neurons (α: Treatment, F(1,20) = 40.4, *p* < 0.001; β: Treatment, F(1,20) = 24.1, *p* < 0.001; Sex by Treatment, F(1,20) = 20.8, *p* < 0.001). The proportion of phosphorylated GSK-3 was greater in VPA neurons, indicative of overall lower activity (Fig. 5C, right panel; α: Treatment, F(1,20) = 21.8, *p* < 0.001; β: Treatment, F(1,20) = 13.0, *p* = 0.002).

**Figure 5.**
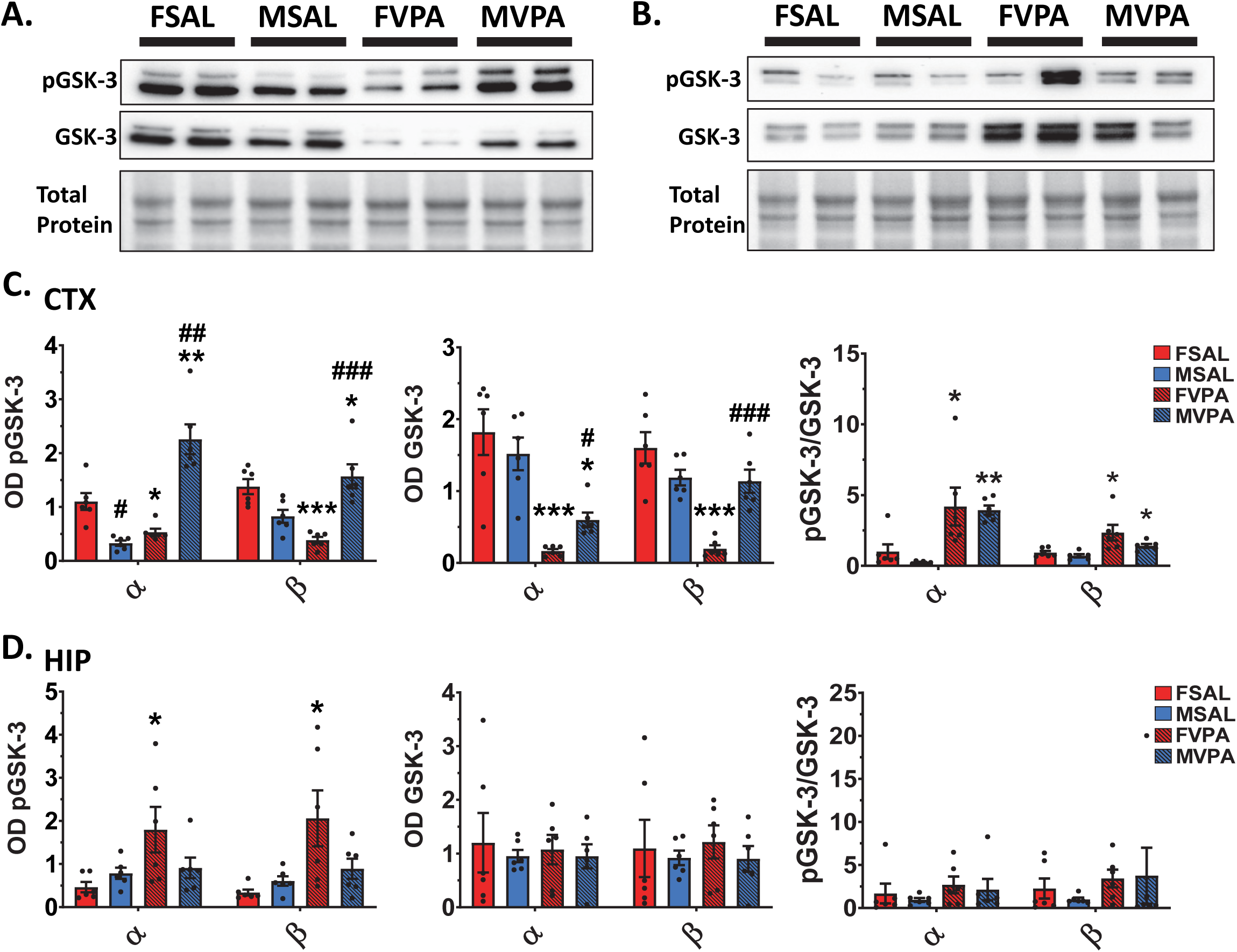
VPA-induced suppression of CTX GSK-3 activity. **A, B)** Representative blots from CTX and HIP neurons showing expression of each isoform of pGSK-3 and GSK-3. **C)** Optical density for CTX pGSK-3 (left panel), total GSK-3 (center panel), and the ratio of pGSK-3 to total GSK-3 (right panel). **D)** Optical density for HIP pGSK-3 (left panel), total GSK-3 (center panel), and the ratio of pGSK-3 to total GSK-3 (right panel). Total protein was used as a loading control. Data are expressed as means ± SEM. N = 6 biological replicates. * *p* < 0.05, ** *p* < 0.01, *** *p* < 0.001 compared to sex-matched controls, ^#^ *p* < 0.05, ^##^ *p* < 0.01, ^###^ *p* < 0.001 compared to females of the same model, ANOVA followed by Bonferroni or Games–Howell post-hoc determined by Levene’s test of variance.

FVPA HIP neurons had higher levels of pGSK-3α (*p* = 0.026) and pGSK-3β (*p* = 0.012) compared to FSAL neurons, while no treatment differences were observed in the male-derived HIP neurons (Fig. 5D, left panel; α: Treatment, F(1,20) = 5.7, *p* = 0.026; β: Treatment, F(1,20) = 8.2, *p* = 0.01). No significant differences were observed in HIP neurons in total or ratio of expression of GSK-3α or GSK-3β (Fig. 5D, center and right panels).

### Sex-specific VPA-induced alterations in CTX gene expression

The most robust sex differences in neuronal structure and function were observed in the VPA-derived CTX neurons. Therefore, VPA-induced alterations in CTX gene expression in the tissue of male and female rat pups at postnatal day 0 were evaluated. There were 176 genes for females, and 149 genes for males that showed a ±1.3-fold change (log_2_ fold change ≤ −0.11 or ≥ 0.11) in expression (*p* < 0.01), with 37 genes overlapping (Fig. 6A), 11 of which were unknown. Overall, FVPA CTX tissue had a greater downregulation of genes (72%) compared to FSAL tissue, while MVPA CTX had a greater upregulation of genes (77%) compared to their controls (Fig. 6B). The 37 genes shared between the FVPA and MVPA rats and their encoded protein function are described in Supplemental Table 2. Volcano plots are displayed in Figure 6C and 6D. Gene ontology identified sex-specific hit terms, with the top four from each group (biological processes, cellular components, and molecular function) shown for females and males in Figure 6E and 6F, respectively. FVPA rats displayed “ion binding”, “extracellular region” and “response to organic substance” as top terms, whereas MVPA rats had “receptor binding”, “neuron part/projection” and “regulation of signaling” as top terms (Fig. 6E, F). The expression of genes involved in growth factor signaling, such as *egfr*, *fgf19,* and *hgf,* were elevated in FVPA CTX (Fig. 6G). Conversely, neuropeptide genes such as *tac1, cartpt, pdyn,* and *penk* were downregulated in the FVPA CTX, as were many receptor genes including *drd1a*, *drd2*, *adora2a*, *mc4r*, and *htr3a* (Fig. 6H). In contrast, MVPA CTX showed elevations in neuropeptide gene expression of *tac1, cartpt, pdyn,* and *penk* (Fig. 6H) as well as in receptors such as *drd1a*, *adora2a, and mc4r* (Fig. 6I).

**Figure 6.**
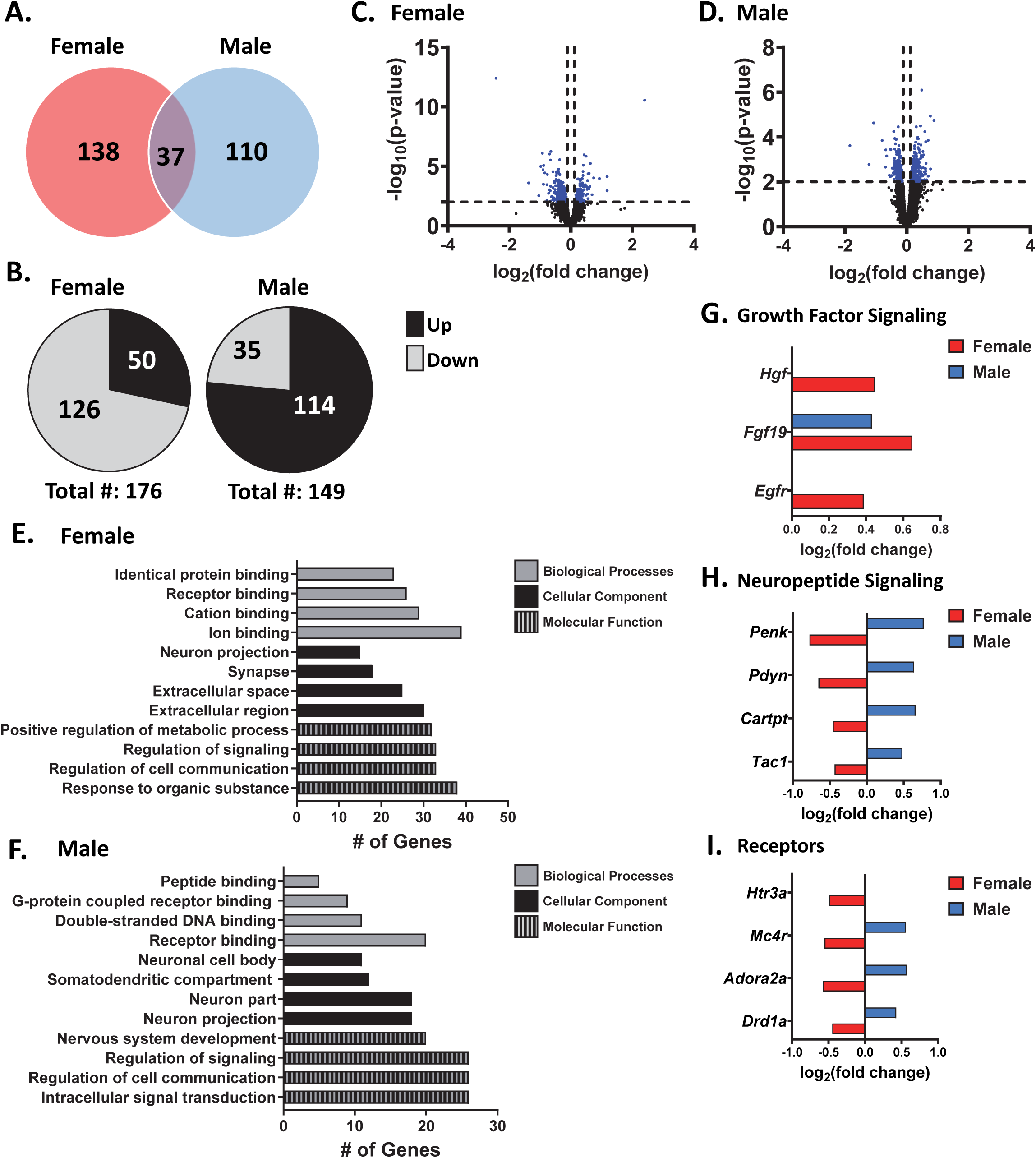
Prenatal VPA induces sex-specific changes in CTX gene expression in rats at birth. **A)** The Venn diagram depicts the number of CTX transcripts that have a ±1.3-fold change for female (red) and male (blue) of VPA tissue **B)** Pie charts depicting the directionality of genes for each sex. **C-D)** Volcano plots represent the −log_10_(p-value) against the log_2_(fold change) magnitude of transcripts altered by VPA for each sex. **E-F)** Gene ontology analysis identified significantly enriched terms for CTX tissue from each sex following VPA in the categories biological processes (grey), cellular components (black), and molecular function (black with grey stripes). **G-I)** Differentially expressed genes for each sex following exposure to VPA in growth factor signaling, neuropeptide signaling, and receptors for both females (red) and males (blue). Data were corrected to *p* ≤ 0.01. N = 5 rats/group.

## Discussion

In the present study, we performed an in-depth assessment of sex differences in neuronal systems function, morphology, and GSK-3 signaling in CTX and HIP neurons derived from the VPA rodent model of ASD, and evaluated sex-specific alterations in CTX gene expression *in vivo*. Overall, there were more pronounced VPA-induced morphological and functional changes in the CTX neurons. With respect to sex differences in neuronal activity, various measures indicated an elevation in activity in VPA neurons in both regions, though these effects were more robust in the female neurons. Both sexes showed region-specific disorganization of firing patterns with VPA. Region-specific changes in dendritic complexity were also evident in the VPA-neurons, which had a greater effect on female-derived neurons. Both sexes showed a relative reduction in GSK-3 activity in CTX neurons only. Transcriptomic analysis revealed differential regulation of CTX genes related to receptors and neuropeptide signaling between the sexes. Taken together, these findings demonstrate that environmental exposure to VPA induces sex-specific neurodevelopmental alterations and support the idea that the heterogeneous manifestation of ASD symptoms resulting from environmental perturbations may have distinct sex-specific neurodevelopmental underpinnings.

In humans, some electroencephalography (EEG) and functional magnetic resonance imaging studies have demonstrated sex-specific changes in neural activity in autistic individuals [29]. For example, one notable study with inclusion criteria that considered behavioral characteristics showed that male autistic individuals displayed hypoconnectivity in the default mode network, whereas females displayed hyperconnectivity [44]. The activity dynamics of neurons either cultured from animal models of ASD or derived from human induced pluripotent stem cells (iPSCs) have also shed some light on mechanistic functional neurodevelopmental changes that may contribute to or initiate the alterations in systems communication that are observed *in vivo* [45–47]. Hyperexcitability and increased firing synchrony in cortical neurons derived from iPSCs that underwent a knockout of an autism risk gene *trpc6* (transient receptor potential canonical 6) have been reported [47]. Hyperexcitability was also observed in Fragile X syndrome iPSC derived neurons [46]. Conversely, there was reduced excitability in human iPSC-derived glutamatergic neurons with gene deletion of the autism risk gene *scn2a* (sodium voltage gated channel alpha subunit 2 alpha) [45]. *Ex vivo* brain slice preparations from VPA rats have also demonstrated hyperexcitable circuits though it must be noted that the sex of the animals used in these studies were not disclosed, and likely pooled [48,49]. Our findings are consistent with these studies, with sex-specific effects on neuronal function also evident. While both cortical FVPA and MVPA neurons showed elevated activity, this effect was more robust in the FVPA neurons. Hyperactivity of HIP neurons was also evident but was female-specific. In addition to neuronal activity, neural activity variability has been proposed as a clinical biomarker in ASD through EEG recordings [50]. Our findings in the VPA neurons are consistent with this research, whereby VPA induced more disorganized firing in both male and female CTX neurons. In the HIP, opposing effects of VPA were observed, with FVPA neurons showing greater organization while MVPA neurons were more disorganized. Together, these findings indicate that there is model- and region-specific alterations in neuronal systems function and that sex plays an important role in these alterations.

Past research has shown that width [51] and length of dendrites [52], number of dendritic spines [53], and AIS length [54–56] can influence neural activity through their effects on protein dynamics, including alterations to populations of voltage-gated sodium and potassium channels. Moreover, altered soma volume may affect the composition of cytoplasmic machinery and protein activity to alter electrical output in ASD [57]. Throughout neuron growth, the proper development of neurons is required to create functional neuronal circuits, which are also said to be dynamic depending on the activity of the circuits [58]. Ghiretti and Paradis (2014) have discussed the adaptability of neurons to become more complex through dendritic arborization as electrical activity at a neuron increases, and can create changes to dendritic morphology and spine number [59]. Post-mortem analyses of children diagnosed with ASD have shown reduced dendritic complexity in HIP CA1 pyramidal neurons [24]. Another study using post-mortem tissue from autistic individuals (age range 10-44) revealed a significant elevation in the dendritic spine number of neurons in layer II pooled from the frontal, temporal, and parietal lobes [60]. In translational studies, there was a reported reduction in dendritic complexity in PFC and HIP neurons *ex vivo* from the C58/J model of ASD [61]. Further, previous work has identified shorter AIS length in the layer II/III neurons from the PFC in *Zbtb16* knockout model of ASD [62], while increased AIS length of CA1 neurons has been reported in the Fragile X model of ASD, which the authors suggest leads to increased excitability [63]. In our findings, CTX FVPA neurons showed less complexity, were shorter in length, and had an elevated number of dendritic spines compared to FSAL neurons. The MVPA neurons were also less complex, though to a lesser degree than the female neurons, and had a reduced soma volume and shorter AIS length compared to controls. In the HIP, FVPA neurons displayed increased complexity, had greater width of dendrites, and more dendritic spines proximally compared to controls, while MVPA neurons displayed a moderate increase in complexity. Our findings presented here are in line with much of the past research from other models; however, there are some differences, reinforcing that heterogeneity in ASD and the type of model employed, as well as the sex of the subject, have a strong influence on the observed neuronal structural alterations.

GSK-3 is a serine/threonine protein kinase highly expressed in the central nervous system. There are two isoforms, alpha (α) and beta (β), with the β isoform having been more extensively researched due to its involvement in numerous cell functions, including synaptic plasticity, signaling, and neuron growth [64–66]. Likely related to those functions, GSK-3 also influences neuronal activity. Indeed, the viral-mediated upregulation of GSK-3β in the prefrontal cortex of rats was shown to dysregulate neuronal oscillatory activity and communication within and between the prefrontal cortex and ventral HIP [67]. Given its widespread role in the CNS, it is not surprising that GSK-3 has been implicated in numerous neurological and neuropsychiatric disorders, including Alzheimer’s disease, Parkinson’s disease, and schizophrenia [68,69]. Variations in the expression and activity of GSK-3β in ASD have also been reported, albeit with conflicting findings. Onore and colleagues isolated T cells from the peripheral blood of adolescent children diagnosed with ASD [70]. They showed increased phosphorylated GSK-3α and GSK-3β levels, indicating reduced activity, with no change in total GSK-3 levels [70]. In animal studies modeling ASD, increased [27,71] or decreased [16,28,72] activity of GSK-3β has been reported. For example, in the Fragile X model of ASD, GSK-3α and GSK-3β activity is elevated in the striatum, CTX, and HIP [27,71]. Conversely, suppressed GSK-3β activity has been previously reported in the CTX, HIP, cerebellum [28], and amygdala [72] of VPA rodents and in the anterior cingulate cortex of *shank3b* knockout mice [16], in agreement with the findings presented in the present study.

A study by Peineau et al. showed that long-term potentiation induced by tetanic stimulation *ex vivo* reduced the activity of GSK-3β, which was proposed to lead to synaptogenesis [66]. Given these findings, it is possible that there was a relationship between the lower GSK-3β expression and activity levels observed in CTX FVPA neurons observed in the present study and the associated elevation in dendritic spines, neither of which were observed in the MVPA CTX neurons. However, both FVPA and MVPA CTX neurons showed relative suppression in CTX GSK-3β activity. Evidence indicates an inhibitory role for GSK-3β on neuronal excitability through its regulation of ion channels [73]. This supports the idea of a link between the more robust alterations in GSK-3β and the greater elevation in neuronal activity observed in the CTX FVPA neurons. However, given that HIP FVPA neuronal activity was also elevated and reduced expression and/or activity GSK-3 was not evident in this region, this indicates that there are likely many more factors contributing to neuronal activity alterations that were not elucidated in the present study. It is worth noting that GSK-3α activity was also suppressed in the CTX VPA neurons of both sexes. GSK-3α is understudied in the context of neurodevelopmental disorders, though the two isoforms have been suggested to be involved in variable cellular functions, such as the conversion from radial to intermediate progenitor cells [74]. Overall, GSK-3 inhibition has previously been proposed as a therapeutic target in regard to ASD [75–77], however much more research is required to elucidate under which conditions this may be appropriate and how sex may influence treatment response.

The most robust alterations in cortical gene expression were associated with the same processes in newborn males and females, namely growth factors, and neuropeptide and receptor signaling. However, aside from *fgf19*, which was upregulated in both sexes, prenatal VPA exposure induced opposing effects on gene expression. This provides additional evidence that the neuronal structural and functional alterations reported herein may involve discrete sex-dependant neurodevelopmental mechanistic underpinnings. Dopamine system dysfunction in ASD has been widely studied, as reviewed by Pavăl [78], and a dopamine transporter knockout animal model has even been proposed to recapitulate ASD symptoms, as it displays social deficits in the social hierarchy and tube dominance test [79]. Alterations in dopamine transmission may, therefore, play a role in the observed changes in neuronal structure and/or function reported herein, though it is important to note here that there exists as well robust sex differences in dopaminergic functioning [80]. Our findings also showed alterations in the gene expression of the endogenous opioids dynorphin and enkephalin, and there is some evidence of alterations of endogenous opioids being altered in autistic children [81]. Opioid antagonists have also been assessed for behaviors related to self-injury and reduced empathy [81,82]. In addition, agonism of the melanocortin receptor 4 has been suggested as a potential therapeutic intervention for ASD due to the receptor concentration on oxytocin-expressing neurons [83], though to normalize dopamine dysfunction potentially. Although, based off the findings presented here, this may only pose a benefit for FVPA rodents. The *adora2a* gene has also been studied in the context of ASD with specific variants associated with increased anxiety [84]. Only the FVPA animals displayed a change in the *htr3a* gene having lower expression, and *htr3a* knockout mice, another proposed model of ASD, exhibit reduced social interaction and increased repetitive behaviour [85].

### Limitations

The present study is a characterization of changes in the VPA rodent model, so caution should be taken when extrapolating the findings to other models. Most of the study was also conducted in cultured primary neurons, which are not necessarily representative of the *in vivo* functional system. These studies were also conducted before behavioral symptoms would normally emerge. Thus, while these findings provide insights into sex-specific neurodevelopmental changes with prenatal VPA exposure, they may not correlate or be causal to the behavioral alterations that occur later during childhood and adolescence.

### Conclusions

In conclusion, we provide evidence that there are region- and sex-specific alterations in neuronal structure and function induced by prenatal exposure to VPA, the most widely used model to study the role of the environment in idiopathic ASD. While this work also offers further validation for using the model when studying specific aspects of the disorder, it is also a call to action to incorporate both sexes in basic, translational, and clinical research. The lack of inclusion of both sexes would hinder diagnosis and future treatment approaches for those who require it. In line with clinical findings that show sex-specific differences in prevalence, age of onset, and symptoms in ASD, this work reinforces that the mechanisms that may drive these differences are most likely different, and thus, more targeted therapies need to be developed.

## Data availability

The datasets used and/or analyzed during the current study are available at the OSF repository osf.io/jy3x9 with gene expression data also available on the Gene Expression Omnibus.

## Acknowledgements

We would like to acknowledge the land in Ontario, Canada, on which this research was performed, the ancestral lands of the Attawandaron people, and the treaty lands and territory of the Mississaugas of the Credit First Nation. We also offer our respect to all of the First Nations, Inuit, and Métis peoples, acknowledging their spirituality, traditional knowledge, and cultural diversity. We offer our gratitude for their environmental stewardship from time immemorial.

## Funding

This work was supported by an award from The Scottish Rite Charitable Foundation of Canada and the Natural Sciences and Engineering Research Council of Canada (to MLP).

## List of Abbreviations

Α: alpha
AIS: axon initial segment
ANOVA: analysis of variance
AraC: Beta-D-arabinofuranoside
ASD: autism spectrum disorder
Β: beta
BSA: bovine serum albumin
CTX: cortical
CV: coefficient of variation
DIV: days in vitro
ECL: enhanced chemiluminescence
EEG: electroencephalography
FSAL: female saline
FVPA: female VPA
GABA: gamma-aminobutyric acid
GO: gene ontology
GSK-3: glycogen synthase kinase 3
HIP: hippocampal
HRP: horseradish peroxidase
ICC: immunocytochemistry
i.p.: intraperitoneal
iPSC: induced pluripotent stem cells
ISI: inter-spike interval
MAP2: microtubule associated protein 2
MEA: multi-electrode array
MFR: mean firing rate
MSAL: male saline
MVPA: male VPA
NGS: normal goat serum
PBS: phosphate-buffered saline
PCA: principal components analysis
PFA: paraformaldehyde
PVDF: polyvinylidene difluoride
RFU: raw fluorescence units
RNA: ribonucleic acid
SDS: sodium dodecyl sulfate
SPSS: statistical package for the social sciences
TBS: tris-buffered saline
TBS-T: tris-buffered saline with tween
VPA: valproic acid

## Notes

https://www.ncbi.nlm.nih.gov/geo/query/acc.cgi?acc=GSE275787

### Competing Interest Statement

The authors have declared no competing interest.

## References

1. American Psychiatric Association. Diagnostic and statistical manual of mental disorders. Fifth Edition. 2013 [cited 2024 May 30]. Available from: https://dsm.psychiatryonline.org/doi/book/10.1176/appi.books.9780890425596

2. Masi A, DeMayo MM, Glozier N, Guastella AJ. An overview of autism spectrum disorder, heterogeneity and treatment options. Neurosci Bull. 2017;33:183–193.

3. Carter AS, Black DO, Tewani S, Connolly CE, Kadlec MB, Tager-Flusberg H. Sex differences in toddlers with autism spectrum disorders. J Autism Dev Disord. 2007;37:86–97.

4. Alaghband-rad J, Hajikarim-Hamedani A, Motamed M. Camouflage and masking behavior in adult autism. Front Psychiatry. 2023;14:1108110.

5. Andrews DS, Diers K, Lee JK, Harvey DJ, Heath B, Cordero D, et al. Sex differences in trajectories of cortical development in autistic children from 2–13 years of age. Mol Psychiatry. 2024;1–12.

6. Frazier TW, Georgiades S, Bishop SL, Hardan AY. Behavioral and cognitive characteristics of females and males with autism in the simons simplex collection. J Am Acad Child Adolesc Psychiatry. 2014;53:329–340.e1-3.

7. Mandy W, Chilvers R, Chowdhury U, Salter G, Seigal A, Skuse D. Sex differences in autism spectrum disorder: evidence from a large sample of children and adolescents. J Autism Dev Disord. 2012;42:1304–1313.

8. Rutherford M, McKenzie K, Johnson T, Catchpole C, O’Hare A, McClure I, et al. Gender ratio in a clinical population sample, age of diagnosis and duration of assessment in children and adults with autism spectrum disorder. Autism. 2016;20:628–634.

9. Banach R, Thompson A, Szatmari P, Goldberg J, Tuff L, Zwaigenbaum L, et al. Brief report: relationship between non-verbal IQ and gender in autism. J Autism Dev Disord. 2009;39:188– 193.

10. Fombonne E. Epidemiological surveys of autism and other pervasive developmental disorders: an update. J Autism Dev Disord. 2003;33:365–382.

11. Khalifa D, Shahin O, Salem D, Raafat O. Serum glutamate was elevated in children aged 3-10 years with autism spectrum disorders when they were compared with controls. Acta Paediatr. 2019;108:295–299.

12. Shinohe A, Hashimoto K, Nakamura K, Tsujii M, Iwata Y, Tsuchiya KJ, et al. Increased serum levels of glutamate in adult patients with autism. Prog Neuropsychopharmacol Biol Psychiatry. 2006;30:1472–1477.

13. Dhossche D, Applegate H, Abraham A, Maertens P, Bland L, Bencsath A, et al. Elevated plasma gamma-aminobutyric acid (GABA) levels in autistic youngsters: stimulus for a GABA hypothesis of autism. Med Sci Monit. 2002;8:PR1-6.

14. El-Ansary A, Al-Ayadhi L. GABAergic/glutamatergic imbalance relative to excessive neuroinflammation in autism spectrum disorders. J Neuroinflammation. 2014;11:189.

15. Yip J, Soghomonian J-J, Blatt GJ. Increased GAD67 mRNA expression in cerebellar interneurons in autism: implications for purkinje cell dysfunction. Journal of Neuroscience Research. 2008;86:525–530.

16. Wang M, Liu X, Hou Y, Zhang H, Kang J, Wang F, et al. Decrease of GSK-3β activity in the anterior cingulate cortex of shank3b−/− mice contributes to synaptic and social deficiency. Frontiers in Cellular Neuroscience. 2019;13:447.

17. Carlsson ML. Hypothesis: is infantile autism a hypoglutamatergic disorder? Relevance of glutamate – serotonin interactions for pharmacotherapy. J Neural Transm. 1998;105:525–535.

18. Horder J, Petrinovic MM, Mendez MA, Bruns A, Takumi T, Spooren W, et al. Glutamate and GABA in autism spectrum disorder—a translational magnetic resonance spectroscopy study in man and rodent models. Transl Psychiatry. 2018;8:106.

19. Robertson CE, Ratai E-M, Kanwisher N. Reduced GABAergic action in the autistic Brain. Current Biology. 2016;26:80–85.

20. Berry-Kravis E, Sumis A, Hervey C, Nelson M, Porges SW, Weng N, et al. Open-label treatment trial of lithium to target the underlying defect in fragile x syndrome. Journal of Developmental & Behavioral Pediatrics. 2008;29:293–302.

21. Porceddu PF, Ciampoli M, Romeo E, Garrone B, Durando L, Milanese C, et al. The novel potent GSK3 inhibitor AF3581 reverts fragile X syndrome phenotype. Human Molecular Genetics. 2022;31:839–849.

22. Bringas ME, Carvajal-Flores FN, López-Ramírez TA, Atzori M, Flores G. Rearrangement of the dendritic morphology in limbic regions and altered exploratory behavior in a rat model of autism spectrum disorder. Neuroscience. 2013;241:170–187.

23. Cloarec R, Riffault B, Dufour A, Rabiei H, Gouty-Colomer L-A, Dumon C, et al. Pyramidal neuron growth and increased hippocampal volume during labor and birth in autism. Science Advances. 2019;5:eaav0394.

24. Raymond GV, Bauman ML, Kemper TL. Hippocampus in autism: a golgi analysis. Acta Neuropathol. 1995;91:117–119.

25. Snow WM, Hartle K, Ivanco TL. Altered morphology of motor cortex neurons in the VPA rat model of autism. Dev Psychobiol. 2008;50:633–639.

26. Sosa-Díaz N, Bringas ME, Atzori M, Flores G. Prefrontal cortex, hippocampus, and basolateral amygdala plasticity in a rat model of autism spectrum. Synapse. 2014;68:468–473.

27. Min WW, Yuskaitis CJ, Yan Q, Sikorski C, Chen S, Jope RS, et al. Elevated glycogen synthase kinase-3 activity in fragile x mice: key metabolic regulator with evidence for treatment potential. Neuropharmacology. 2009;56:463–472.

28. Qin L, Dai X, Yin Y. Valproic acid exposure sequentially activates wnt and mTOR pathways in rats. Mol Cell Neurosci. 2016;75:27–35.

29. Williams OOF, Coppolino M, Perreault ML. Sex differences in neuronal systems function and behaviour: beyond a single diagnosis in autism spectrum disorders. Transl Psychiatry. 2021;11:625.

30. Perreault ML. Consideration of Research Approaches in Systems Neurobiology. Biol Psychiatry Glob Open Sci. 2024;4:107–109.

31. Uhlhaas PJ, Roux F, Rodriguez E, Rotarska-Jagiela A, Singer W. Neural synchrony and the development of cortical networks. Trends Cogn Sci. 2010;14:72–80.

32. Zikopoulos B, Barbas H. Altered neural connectivity in excitatory and inhibitory cortical circuits in autism. Frontiers in Human Neuroscience. 2013;7:609.

33. Thériault R-K, Manduca JD, Perreault ML. Sex differences in innate and adaptive neural oscillatory patterns link resilience and susceptibility to chronic stress in rats. J Psychiatry Neurosci. 2021;46:E258–E270.

34. Nicolini C, Fahnestock M. The valproic acid-induced rodent model of autism. Experimental Neurology. 2018;299:217–227.

35. Gouda B, Sinha SN, Chalamaiah M, Vakdevi V, Shashikala P, Veeresh B, et al. Sex differences in animal models of sodium-valproate-induced autism in postnatal BALB/c mice: whole-brain histoarchitecture and 5-HT2A receptor biomarker evidence. Biology. 2022;11:79.

36. Kazlauskas N, Seiffe A, Campolongo M, Zappala C, Depino AM. Sex-specific effects of prenatal valproic acid exposure on sociability and neuroinflammation: relevance for susceptibility and resilience in autism. Psychoneuroendocrinology. 2019;110:104441.

37. Melancia F, Schiavi S, Servadio M, Cartocci V, Campolongo P, Palmery M, et al. Sex-specific autistic endophenotypes induced by prenatal exposure to valproic acid involve anandamide signalling. British Journal of Pharmacology. 2018;175:3699–3712.

38. Thériault R-K, St-Denis M, Hewitt T, Khokhar JY, Lalonde J, Perreault ML. Sex-specific cannabidiol- and iloperidone-induced neuronal activity changes in an in vitro MAM model system of schizophrenia. Int J Mol Sci. 2021;22:5511.

39. Oorschot DE, Jones DG. Non-neuronal cell proliferation in tissue culture: implications for axonal regeneration in the central nervous system. Brain Research. 1986;368:49–61.

40. McCready FP, Gordillo-Sampedro S, Pradeepan K, Martinez-Trujillo J, Ellis J. Multielectrode arrays for functional phenotyping of neurons from induced pluripotent stem cell models of neurodevelopmental disorders. Biology (Basel). 2022;11:316.

41. Hasbi A, Fan T, Alijaniaram M, Nguyen T, Perreault ML, O’Dowd BF, et al. Calcium signaling cascade links dopamine D1–D2 receptor heteromer to striatal BDNF production and neuronal growth. Proc Natl Acad Sci U S A. 2009;106:21377–21382.

42. Perreault ML, Jones-Tabah J, O’Dowd BF, George SR. A physiological role for the dopamine D5 receptor as a regulator of BDNF and Akt signalling in rodent prefrontal cortex. International Journal of Neuropsychopharmacology. 2013;16:477–483.

43. McCallum RT, Thériault R-K, Manduca JD, Russell ISB, Culmer AM, Doost JS, et al. Correction: Nrf2 activation rescues stress-induced depression-like behaviour and inflammatory responses in male but not female rats. Biol Sex Differ. 2024;15:31.

44. Alaerts K, Swinnen SP, Wenderoth N. Sex differences in autism: a resting-state fMRI investigation of functional brain connectivity in males and females. Social Cognitive and Affective Neuroscience. 2016;11:1002–1016.

45. Brown CO, Uy JA, Murtaza N, Rosa E, Alfonso A, Dave BM, et al. Disruption of the autism-associated gene SCN2A alters synaptic development and neuronal signaling in patient iPSC- glutamatergic neurons. Front Cell Neurosci. 2024;17:1239069.

46. Shen M, Sirois CL, Guo Y, Li M, Dong Q, Méndez-Albelo NM, et al. Species-specific FMRP regulation of RACK1 is critical for prenatal cortical development. Neuron. 2023;43:114330

47. Shin KC, Ali G, Ali Moussa HY, Gupta V, de la Fuente A, Kim H-G, et al. Deletion of TRPC6, an autism risk gene, induces hyperexcitability in cortical neurons derived from human pluripotent stem cells. Mol Neurobiol. 2023;60:7297–7308.

48. Lin H-C, Gean P-W, Wang C-C, Chan Y-H, Chen PS. The amygdala excitatory/inhibitory balance in a valproate-induced rat autism model. PLos One. 2013;8:e55248.

49. Rinaldi T, Kulangara K, Antoniello K, Markram H. Elevated NMDA receptor levels and enhanced postsynaptic long-term potentiation induced by prenatal exposure to valproic acid. Proc Natl Acad Sci USA. 2007;104:13501–13506.

50. Hecker L, Wilson M, Tebartz van Elst L, Kornmeier J. Altered EEG variability on different time scales in participants with autism spectrum disorder: an exploratory study. Sci Rep. 2022;12:13068.

51. Alcami P, El Hady A. Axonal computations. Front Cell Neurosci. 2019; 13:413.

52. Ilmoniemi RJ, Mäki H, Saari J, Salvador R, Miranda PC. The frequency-dependent neuronal length constant in transcranial magnetic stimulation. Front Cell Neurosci. 2016;10:194.

53. Aizenman CD, Huang EJ, Linden DJ. Morphological correlates of intrinsic electrical excitability in neurons of the deep cerebellar nuclei. J Neurophysiol. 2003;89:1738–1747.

54. Bonnevie VS, Dimintiyanova KP, Hedegaard A, Lehnhoff J, Grøndahl L, Moldovan M, et al. Shorter axon initial segments do not cause repetitive firing impairments in the adult presymptomatic G127X SOD-1 amyotrophic lateral sclerosis mouse. Sci Rep. 2020;10:1280.

55. Salzer, J. L. An unfolding role for ankyrin-G at the axon initial segment. Proc Natl Acad Sci. 2019;116:19228–19230.

56. Leterrier C. The axon initial segment: an updated viewpoint. J Neurosci. 2018;38:2135– 2145.

57. Wegiel J, Flory M, Schanen NC, Cook EH, Nowicki K, Kuchna I, et al. Significant neuronal soma volume deficit in the limbic system in subjects with 15q11.2-q13 duplications. Acta Neuropathol Commun. 2015;3:63.

58. Forrest MP, Parnell E, Penzes P. Dendritic structural plasticity and neuropsychiatric disease. Nat Rev Neurosci. 2018;19:215–234.

59. Ghiretti AE, Paradis S. Molecular mechanisms of activity-dependent changes in dendritic morphology: role of RGK proteins. Trends Neurosci. 2014;37:399–407.

60. Hutsler JJ, Zhang H. Increased dendritic spine densities on cortical projection neurons in autism spectrum disorders. Brain Research. 2010;1309:83–94.

61. Barón-Mendoza I, Maqueda-Martínez E, Martínez-Marcial M, De la Fuente-Granada M, Gómez-Chavarin M, González-Arenas A. Changes in the number and morphology of dendritic spines in the hippocampus and prefrontal cortex of the C58/J mouse model of autism. Front Cell Neurosci. 2021;15:726501.

62. Usui N, Tian X, Harigai W, Togawa S, Utsunomiya R, Doi T, et al. Length impairments of the axon initial segment in rodent models of attention-deficit hyperactivity disorder and autism spectrum disorder. Neurochemistry International. 2022;153:105273.

63. Booker SA, Oliveira LS de, Anstey NJ, Kozic Z, Dando OR, Jackson AD, et al. Input-output relationship of CA1 pyramidal neurons reveals intact homeostatic mechanisms in a mouse model of fragile x syndrome. Cell Reports. 2020;32:107988.

64. Hur E-M, Zhou F-Q. GSK3 signalling in neural development. Nat Rev Neurosci. 2010;11:539–551.

65. López-Tobón A, Villa CE, Cheroni C, Trattaro S, Caporale N, Conforti P, et al. Human cortical organoids expose a differential function of GSK3 on cortical neurogenesis. Stem Cell Reports. 2019;13:847–861.

66. Peineau S, Taghibiglou C, Bradley C, Wong TP, Liu L, Lu J, et al. LTP Inhibits LTD in the hippocampus via regulation of GSK3β. Neuron. 2007;53:703–717.

67. Albeely AM, Williams OOF, Perreault ML. GSK-3β disrupts neuronal oscillatory function to inhibit learning and memory in male rats. Cell Mol Neurobiol. 2021;42:1341–1353.

68. Albeely AM, Ryan SD, Perreault ML. Pathogenic feed-forward mechanisms in alzheimer’s and parkinson’s disease converge on GSK-3. Brain Plast. 2018;4:151–167.

69. Manduca JD, Thériault RK, Perreault ML. Glycogen synthase kinase-3: the missing link to aberrant circuit function in disorders of cognitive dysfunction? Pharmacol Res. 2020;157:104819.

70. Onore C, Yang H, de Water JV, Ashwood P. Dynamic akt/mTOR signaling in children with autism spectrum disorder. Front Pediatr. 2017;5:43.

71. Yuskaitis CJ, Mines MA, King MK, David Sweatt J, Miller CA, Jope RS. Lithium ameliorates altered glycogen synthase kinase-3 and behavior in a mouse model of fragile x syndrome. Biochem pharmacol. 2010;79:632–646.

72. Wu HF, Chen PS, Chen Y-J, Lee C-W, Chen I-T, Lin H-C. Alleviation of n-methyl-d-aspartate receptor-dependent long-term depression via regulation of the glycogen synthase kinase-3β pathway in the amygdala of a valproic acid-induced animal model of autism. Mol Neurobiol. 2017;54:5264–5276.

73. Jaworski T. Control of neuronal excitability by GSK-3beta: epilepsy and beyond. Biochim Biophys Acta Mol Cell Res. 2020;1867:118745.

74. Ma Y, Wang X, Chen J, Li B, Hur E-M, Saijilafu. Differential roles of glycogen synthase kinase 3 subtypes alpha and beta in cortical development. Front Mol Neurosci. 2017;10:391.

75. Beurel E, Grieco SF, Jope RS. Glycogen synthase kinase-3 (GSK3): regulation, actions, and diseases. Pharmacol Ther. 2015;0:114–131.

76. Mines MA, Jope RS. Glycogen synthase kinase-3: a promising therapeutic target for fragile x syndrome. Front Mol Neurosci. 2011;4:35.

77. Ruiz SMA, Eldar-Finkelman H. Glycogen synthase kinase-3 inhibitors: preclinical and clinical focus on CNS-a decade onward. Front Mol Neurosci. 2022;14:792364.

78. Pavăl D. A dopamine hypothesis of autism spectrum disorder. Dev Neurosci. 2017;39:355– 360.

79. Rodriguiz RM, Chu R, Caron MG, Wetsel WC. Aberrant responses in social interaction of dopamine transporter knockout mice. Behav Brain Res. 2004;148:185–198.

80. Williams OOF, Coppolino M, George SR, Perreault ML. Sex differences in dopamine receptors and relevance to neuropsychiatric disorders. Brain Sci. 2021;11:1199.

81. Gillberg C. Endogenous opioids and opiate antagonists in autism: brief review of empirical findings and implications for clinicians. Dev Med Child Neurol. 1995;37:239–245.

82. Chabane N, Leboyer M, Mouren-Simeoni MC. Opiate antagonists in children and adolescents. Eur Child Adolesc Psychiatry. 2000;9:S44–S50.

83. Minakova E, Lang J, Medel-Matus J-S, Gould GG, Reynolds A, Shin D, et al. Melanotan-II reverses autistic features in a maternal immune activation mouse model of autism. PLoS One. 2019;14:e0210389.

84. Freitag CM, Agelopoulos K, Huy E, Rothermundt M, Krakowitzky P, Meyer J, et al. Adenosine A2A receptor gene (ADORA2A) variants may increase autistic symptoms and anxiety in autism spectrum disorder. Eur Child Adolesc Psychiatry. 2010;19:67–74.

85. Huang L, Wang J, Liang G, Gao Y, Jin S-Y, Hu J, et al. Upregulated NMDAR-mediated GABAergic transmission underlies autistic-like deficits in Htr3a knockout mice. Theranostics. 2021;11:9296–9310.

