## Supplemental Tables for "Prenatal exposure to valproic acid induces sex-specific alterations in cortical and hippocampal neuronal structure and function in rats"

**Supplemental Table 1. ANOVA-derived F-statistics for group comparisons in neuronal activity taken from CTX and HIP neurons.**

| | Sex | Treatment | Sex*Treatment |
|---|---|---|---|
| CTX | | | |
| MFR | F(1,751)=6.5, p=0.011 | F(1,751)=82.5, p<0.001 | F(1,751)=37.3, p<0.001 |
| # of Spikes | F(1,869)=9.4, p=0.002 | F(1,869)=40.6, p<0.001 | F(1,869)=22.7, p<0.001 |
| # of Bursts | F(1,754)=7.9, p=0.005 | F(1,754)=80.0, p<0.001 | F(1,754)=37.6, p<0.001 |
| # of Spikes/Burst | F(1,870)=1.5, p=0.221 | F(1,870)=14.5, p<0.001 | F(1,870)=0.8, p=0.366 |
| Synchrony | F(1,62)=6.9, p=0.011 | F(1,62)=58.1, p<0.001 | F(1,62)=2.2, p=0.140 |
| ISI within a burst | F(1,870)=14.8, p<0.001 | F(1,870)=117.1, p<0.001 | F(1,870)=9.4, p=0.002 |
| Coefficient of Variation | F(1,814)=0.7, p=0.416 | F(1,814)=120.9, p<0.001 | F(1,814)=0.4, p=0.548 |
| HIP | | | |
| MFR | F(1,734)=13.8, p<0.001 | F(1,734)=8.0, p=0.005 | F(1,734)=10.8, p=0.001 |
| # of Spikes | F(1,849)=13.9, p<0.001 | F(1,849)=6.0, p=0.014 | F(1,849)=18.4, p<0.001 |
| # of Bursts | F(1,660)=23.1, p<0.001 | F(1,660)=0.0, p=0.958 | F(1,660)=77.3, p<0.001 |
| # of Spikes/Burst | F(1,738)=4.2, p=0.042 | F(1,738)=8.4, p=0.004 | F(1,849)=18.4, p<0.001 |
| Synchrony Index | F(1,65)=11.9, p<0.001 | F(1,65)=0.0, p=0.982 | F(1,65)=8.3, p=0.077 |
| ISI within a burst | F(1,738)=21.2, p<0.001 | F(1,738)=5.6, p=0.018 | F(1,738)=38.2, p<0.001 |
| Coefficient of Variation | F(1,794)=0.1, p=0.801 | F(1,794)=2.9, p=0.086 | F(1,794)=33.3, p<0.001 |

**Supplemental Table 2. Known genes that showed altered expression in female and male CTX tissue at postnatal day 0 and the description of the genes.**

| Gene | Direction Females | Direction Males | Description |
|---|---|---|---|
| *Fgf19* | ↑ | ↑ | Fibroblast growth factor 19; member of fibroblast growth factor family, has cell survival activities important in cell growth, morphogenesis and tissue repair. Suggested to only be present in fetal tissue. |
| *Arrdc3* | ↑ | ↑ | Arrestin domain containing 3; a member of the arrestin family of proteins, regulates G-protein mediated signaling. |
| *Rarb* | ↓ | ↑ | Retinoic acid receptor, beta; binds retinoic acid (active form of vitamin A), which mediates cellular signaling in embryonic morphogenesis, cell growth, and differentiation. A member of the thyroid-steroid hormone receptor superfamily of nuclear transcriptional regulators. |
| *Epcam* | ↓ | ↑ | Epithelial cell adhesion molecule; is a carcinoma-associated antigen and is found in most epithelial cells, also functions as a homotypic calcium-independent cell adhesion molecule. |
| *Drd1* | ↓ | ↑ | Dopamine receptor D1; encodes for D1 subtype of dopamine receptor, is the most abundant dopamine receptor in the CNS. Regulate neuronal growth/development, behavior, and DRD2-mediated events. G-protein coupled receptor stimulates adenylyl cyclase and activates cyclic AMP-dependent protein kinases. |
| *Cartpt* | ↓ | ↑ | CART prepropeptide; preproprotein that is proteolytically processed to generate multiple biologically active peptides. Plays a role in appetite, energy, balance, reward, addiction. Upregulated following administration of drugs such as amphetamines. Ligand of Nr4a1 (highly expressed in striatal regions; dopaminergic), helps determine striatal dopamine levels. |
| *Gpr88* | ↓ | ↑ | G-protein coupled receptor 88; a protein encoded by this gene is a G-protein coupled receptor most exclusively expressed in the striatum. Deficits associated with neuropsychiatric diseases and learning difficulties/speech delay. |
| *Gnal* | ↓ | ↑ | G-protein subunit alpha L; is a stimulatory G protein alpha which mediates odorant signaling in the olfactory epithelium. It couples dopamine type 1 receptors and adenosine A2A receptors which is widely expressed in the CNS. |

| | | | |
|---|---|---|---|
| *Mc4r* | ↓ | ↑ | Melanocortin 4 receptor; is a membrane-bound receptor and member of the melanocortin receptor family. Interacts with adrenocorticotropic hormone and MSH hormones, mediated by G proteins. |
| *Ebf1* | ↓ | ↑ | EBF transcription factor 1; Predicted to be a positive regulator of transcription by RNA polymerase II. Predicted to be located in the nucleus and a part of chromatin. |
| *Adora2a* | ↓ | ↑ | Adenosine A2a receptor; member of G-protein coupled receptor superfamily. This protein, an adenosine receptor of A2A subtype, uses adenosine as the preferred endogenous agonist and preferentially interacts with the G(s) and G(olf) family of G proteins to increase intracellular cAMP. Involved in several processes (cerebral blood flow, pain regulation, immune function, etc.). |
| *Lamp5* | ↓ | ↑ | Lysosomal-associated membrane protein family, member 5; predicted to be involved in protein localization to organelle. Located in the endoplasmic reticulum of many cell components: Golgi intermediate compartment membrane, endosome membrane and plasma membrane. |
| *Pdyn* | ↓ | ↑ | Prodynorphin; a preproprotein that is proteolytically processed to form secreted opioid peptides: beta-neoendorphin, dynorphin, leu-enkephalin, rimorphin and leumorphin. Ligands for the kappa-type of opiod receptor. Modulates response to psychoactive compounds (including cocaine). |
| *Penk* | ↓ | ↑ | Proenkephalin; a preproprotein that is cleaved into opioid peptides: met-enkephalin and leu-enkephalin. Stored in synaptic vesicles then released into synapse by binding to mu- or delta-opiod receptors to modulate pain perception. |
| *Rxrg* | ↓ | ↑ | Retinoid X receptor gamma; is a member of the retinoid X receptor family of nuclear receptors which mediates the antiproliferative effects of retinoic acid. |
| *Lhx8* | ↓ | ↑ | LIM homeobox 8; involved in patterning and differentiation of various tissues, including neurons. |
| *C5h9orf85* | ↓ | ↓ | Similar to human chromosome 9 open reading frame 85; a protein coding gene. |
| *Ebpl* | ↓ | ↓ | EBP Like; predicted to enable cholestenol delta-isomerase activity. Predicted to be involved in sterol metabolic processes. Located in the endoplasmic reticulum. |
| *Cpa3* | ↓ | ↓ | Carboxypeptidase A3; a preproprotein which, when cleaved, is released by mast cells for degradation of endogenous protein and venom-associated proteins. May be elevated in asthma patients. |

| | | | |
|---|---|---|---|
| *Fcrl2* | ↓ | ↓ | Fc receptor-like 2; a member of the immunoglobulin receptor family. Is one of many Fc receptor-like glycoproteins clustered on the long arm of chromosome 1. |
| *Mcpt1* | ↓ | ↓ | Mast cell protease 1-like 1; predicted to enable serine-type endopeptidase activity. Predicted to be involved in proteolysis and protein coding. Predicted to be active in the cytoplasm, extracellular space, and intracellular membrane-bounded organelles. |
| *Sdr42e1* | ↓ | ↓ | Short chain dehydrogenase/reductase family 42E, member 1; enables oxidoreductase activity. Predicted to be involved in steroid biosynthetic process. Predicted to be an integral part of the membrane. |
| *S100a9* | ↓ | ↓ | S100 calcium binding protein A9; involved in processes including cell cycle progression and differentiation. S100 proteins are localized in the cytoplasm and/or nucleus of numerous cells. Considered an "alarmin" – released upon tissue damage and stimulate response. Major protein in monocytes |
| *Rnf135* | ↓ | ↓ | Ring finger protein 135; predicted to be involved in protein-protein interactions, and protein-DNA interactions. |
